# Mammalian copper homeostasis requires retromer-dependent recycling of the high-affinity copper transporter 1 (CTR1/SLC31A1)

**DOI:** 10.1101/2020.05.26.117150

**Authors:** Rachel Curnock, Peter J. Cullen

## Abstract

The mammalian cell surface is decorated with a plethora of integral membrane proteins including those required for the transport of micronutrients, such as copper, which are essential to cellular health. The concentration of micronutrients within the cell is tightly regulated to avoid their adverse deficiency and toxicity effects. The sorting and recycling of nutrients transporters within the endo-lysosomal network is recognised as an essential process in regulating nutrient balance. The evolutionarily conserved endosomal sorting complex, retromer, coordinates integral membrane protein recognition and retrieval. Cellular copper homeostasis is regulated primarily by two transporters: the major copper influx transporter copper transporter 1 (CTR1/SLC31A1), which controls the uptake of copper from the extracellular environment and is essential for early embryonic development, and the established retromer cargo, the copper-transporting ATPase, ATP7A. Here, we show that in response to fluctuating extracellular copper the retromer complex controls the delivery of CTR1 to the cell surface. Following copper exposure, CTR1 is endocytosed to prevent excessive copper uptake. We reveal that internalised CTR1 localises on retromer-positive endosomes and in response to decreased extracellular copper retromer controls the recycling of CTR1 back to the cell surface to maintain copper homeostasis. In addition to copper, CTR1 plays a central role in platinum uptake. Significantly, the efficacy of platinum-based cancer drugs has been correlated with CTR1 expression. Consistent with this, we demonstrate that retromer-deficient cells show reduced sensitivity to the platinum-based drug, cisplatin.

## INTRODUCTION

Integral membrane proteins are delivered and removed from the cell surface by means of membrane trafficking pathways (Cullen and Steinberg, 2018). The balance between delivery and internalisation defines the steady-state cell surface abundance of these proteins and provides the means to rapidly alter the abundance of individual proteins and larger groups of functionally related proteins in response to environmental cues. For responses triggered by, for example, metabolic cues and toxic stresses, the endocytic network plays an important role in the adaptive response (Corvera et al., 1986; Jenness and Spatrick, 1986; Jones et al., 2012; Mueller et al., 2015; Tanner and Lienhard, 1987). Comprising a number of intracellular membrane bound compartments, this network sorts internalised cell surface integral membrane proteins between two fates: either degradation within the lysosome or retrieval from this fate for subsequent recycling and reuse at the cell surface (Burd and Cullen, 2014; Cullen and Steinberg, 2018). By deciding the fate of proteins that include signalling receptors, nutrient and solute transporters, cell-cell and cell-matrix adhesion molecules, the endocytic network can alter the functionality of the cell surface during a range of extracellular stimuli. However, our understanding of the role of the endocytic retrieval and recycling pathways in adaptive responses has been hampered by a lack of basic understanding of the machinery that regulates these pathways (Bai et al., 2012; Grant and Donaldson, 2009; McNally and Cullen, 2018).

Several protein complexes have now been identified that control the sorting of internalised integral proteins for retrieval and recycling (Arighi et al., 2004; Derivery et al., 2009; Gomez and Billadeau, 2009; McNally et al., 2017; Phillips-Krawczak et al., 2015; Seaman, 2004; Seaman et al., 1998; Simonetti et al., 2019). These include the mammalian retromer, a stable heterotrimeric complex of VPS35, VPS29 and VPS26 of which two isoforms, VPS26A and VPS26B, are expressed in humans (Arighi et al., 2004; Kerr et al., 2005; Seaman, 2004; Seaman et al., 1998). Retromer functions by scaffolding an endosome-associated multi-protein assembly that co-ordinates sequence-dependent cargo selection with the biogenesis of cargo-enriched transport carriers that delivery the sorted cargo back to the cell surface (Burd and Cullen, 2014; Wang et al., 2018). Important retromer accessory proteins include the cargo adaptor sorting nexin-27 (SNX27) and the actin polymerising FAM21-containing WASH (Wiskott-Aldrich syndrome protein and Scar Homology) complex (Gallon et al., 2014; Gomez and Billadeau, 2009; Harbour et al., 2012; Jia et al., 2012; Steinberg et al., 2013; Temkin et al., 2011). Retromer regulates the endosomal retrieval and recycling of numerous integral membrane proteins that include a number of transporters for the uptake and extrusion of essential but also toxic transition metals such as copper (Steinberg et al., 2013). It is clear retromer is central to regulating cargo traffic through the endosomal network and we recently recognised that retromer itself is tightly regulated in response to extracellular cues (Curnock et al., 2019).

Many retromer-dependent cargoes have been identified (Cullen and Steinberg, 2018). The copper-transporting ATPase, ATP7A is an established retromer cargo and its expression at the cell surface upon copper elevation requires retromer and SNX27 (Steinberg et al., 2013). Cellular copper homeostasis is regulated by two principle transporters: the copper pumping ATP7A and ATP7B which sequesters copper into the *trans*-Golgi-network (TGN) and mobilises copper to the cell periphery, and the structurally unrelated cell surface localised copper transporter 1 (CTR1) which controls the uptake of copper from the extracellular environment (Hartwig et al., 2019; Hung et al., 1997; Lee et al., 2002; Petris et al., 1996). Cellular adaptation to elevated extracellular copper is mediated by membrane trafficking pathways that deliver ATP7A to the cell surface to enhance the extrusion of copper, and the endocytosis of CTR1 which serves to prevent further copper uptake and accumulation (Kim et al., 2008). Copper is an essential micronutrient, vital for development and growth but in excess it is toxic (Madsen and Gitlin, 2007). Understandably, homeostatic copper uptake and distribution is tightly controlled. The adverse effects of copper misbalance manifest in the neurological disorders Menkes and Wilson’s diseases, resulting from copper deficiency and overload, respectively (Daniel et al., 2004). Abnormal copper accumulation has also been linked with neuronal cell death and disease pathology in neurodegenerative disorders such as Parkinson’s disease (PD) (Madsen and Gitlin, 2007). It is now well-established that mutations leading to perturbed retromer function are risk factors for the development of PD (McMillan et al., 2017). In addition to the established role for retromer in ATP7A transport, we previously identified the main copper uptake transporter CTR1 as a putative retromer cargo (Steinberg et al., 2013). Therefore, to fully appreciate whether there is a link between retromer function, copper homeostasis and potentially PD progression, in this study we examined the membrane trafficking of CTR1.

The high-affinity copper transporter CTR1 is evolutionarily conserved mediating the cellular import of copper in eukaryotes from fungi to vertebrates and is the only known mammalian importer of copper (Dancis et al., 1994; Zhou and Gitschier, 1997). CTR1 exists in the plasma membrane as homo-trimer with an extracellular N-terminal region, three transmembrane helices and an intracellular C-terminus (Eisses and Kaplan, 2005; Klomp et al., 2002; Lee et al., 2002; Puig et al., 2002; Ren et al., 2019). Deletion studies in mice have demonstrated that CTR1 is essential for development with early embryonic lethality in homozygous knockout animals. Characterisation of heterozygote mice has revealed a severe deficiency in brain copper concentration indicating a crucial role for CTR1 in copper uptake and distribution in the brain (Kuo et al., 2001; Lee et al., 2001). In addition to the cellular import of copper, CTR1 has also been implicated in the cellular uptake and accumulation of platinum-derived anticancer drugs, such as cisplatin, carboplatin, and oxaliplatin (Holzer et al., 2006). Furthermore, responsiveness to these platinum-based chemotherapeutic agents in cancer patients has been correlated with high levels of CTR1 expression (Song et al., 2004). The occupancy and turnover of CTR1 at the plasma membrane is therefore pertinent to not only cellular copper homeostasis but also the efficacy of platinum-based cancer drugs. In the present study, using RNAi-mediated suppression and CRISPR-Cas9-mediated knockout of VPS35, we have investigated whether the cell surface levels and trafficking of CTR1 in response to changes in extracellular copper concentration are dependent on retromer expression. We find using antibody internalisation in the presence of copper and confocal analysis that internalised CTR1 resides on retromer-positive endosomes. Cell surface biotinylation together with flow cytometry revealed that CTR1 basal cell surface levels and copper concentration-dependent delivery to the cell surface relies on retromer expression. The effects of increasing extracellular copper and exposure to cisplatin, the most commonly used platinum chemotherapy drug, on metabolic activity and cell viability were assessed using colorimetric MTT assays. We demonstrate that retromer is essential for full sensitivity to extracellular copper loading and cisplatin toxicity.

## RESULTS

### Retromer is required for the cell surface localization of CTR1

A multitude of metal ion transporters were previously identified as SNX27-retromer cargo using a global unbiased approach (Steinberg et al., 2013). A distinct subset of these transporters were identified as being independent of the accessory protein SNX27 and solely retromer-dependent. The main mammalian copper importer, CTR1 was one such transporter (Steinberg et al., 2013). To validate this finding, SNX27 and the core component of the retromer complex, VPS35 were efficiently suppressed using siRNA and cell surface biotinylation followed by fluorescence-based quantitative western blotting was used to assess the cell surface abundance of CTR1. A significant reduction in the cell surface levels of CTR1 was observed in cells suppressed for VPS35 but not SNX27 (Figure 1A). Notably, as would be anticipated the cell surface levels of ATP7A, an established retromer and SNX27 cargo (Steinberg et al., 2013) was depleted in cells treated with siRNA targeting VPS35 and SNX27 (Figure 1A). Immunofluorescence analysis revealed a reduction in CTR1 signal intensity at the cell surface in VPS35 suppressed cells (Figure 1B). Furthermore, flow cytometry in VPS35 KO HeLa cells confirmed a significant reduction in the abundance of cell surface CTR1 in the absence of retromer function (Figure 1C).

**Figure 1:**
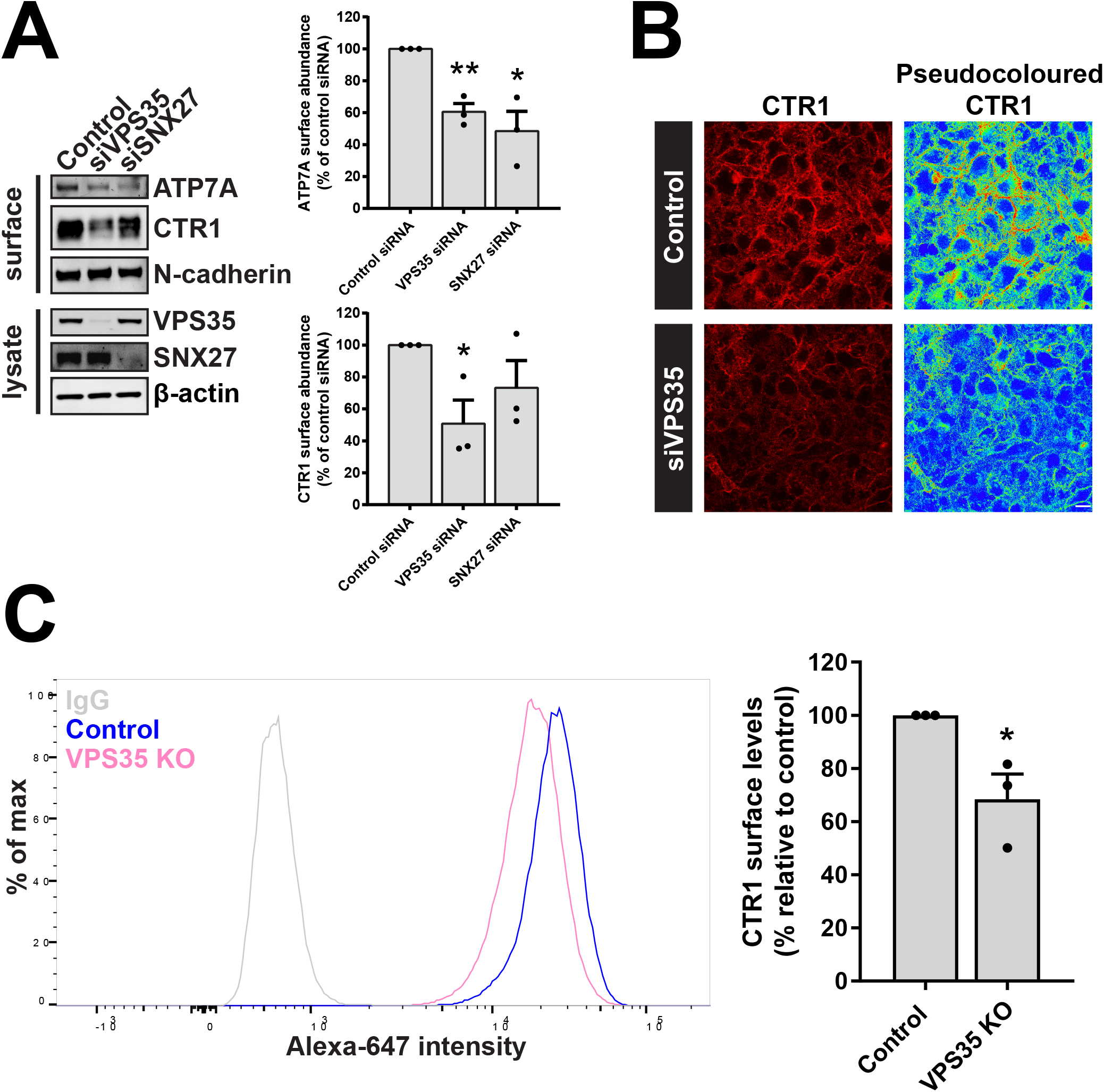
Retromer maintains the cell surface abundance of the evolutionarily conserved copper importer, CTR1. (A) HeLa cells were treated with non-target siRNA or siRNA targeting VPS35 or SNX27 for 72 hours. Cells were then surface biotinylated and streptavidin agarose used to capture biotinylated membrane proteins. Surface abundance of the indicated proteins were assessed by immunoblotting. Data were normalized to N-cadherin. The quantification shows the mean ± SEM; *n* = 3 independent experiments; Student’s *t* test (unpaired). (B) HeLa cells were treated with non-target control siRNA or siRNA targeting VPS35 for 72 hours. Cells were then immunostained for cell surface CTR1 (red). The pseudocoloured panel represents the CTR1 signal intensity. Scale bar, 20 μm. (C) Quantitative analysis by flow cytometry of surface CTR1 levels in control and VPS35 KO HeLa cells. Bars represent the change in CTR1 cell surface levels in VPS35 KO cells versus control cells. Data represent mean ± SEM; *n* = 3 independent experiments; Student’s *t* test (unpaired); \**P* < 0.05; \*\**P* < 0.01.

In response to elevated extracellular copper, CTR1 is internalised in a dynamin- and clathrin-dependent manner to prevent excess copper uptake (Clifford et al., 2016). To visualise the trafficking of CTR1, we performed antibody internalisation in the presence of copper (200 μM CuCl_2_). Here, cells on ice were surface labelled with an antibody against an exofacial epitope of CTR1 and then allowed to internalise the antibody bound CTR1 by incubation at 37°C in growth media supplemented with 200 μM CuCl_2_. Cells were subsequently fixed prior to immunofluorescence and colocalization analysis using LAMP1 as a marker of the degradative lysosomal compartment. After 2 hours in the presence of copper, a significant increase in colocalization between CTR1 and LAMP1 was observed in HeLa cells treated with siRNA targeting VPS35 (Figure 2A) and in VPS35 KO HeLa cells (Figure 2B). Under these conditions, in control cells and following RNAi-mediated suppression or CRISPR-Cas9-mediated knockout of SNX27 CTR1 localised to the cell peripherally or LAMP1-negative intracellular compartments. SNX27 does not therefore make a significant contribution to the copper-dependent sorting of CTR1.

**Figure 2:**
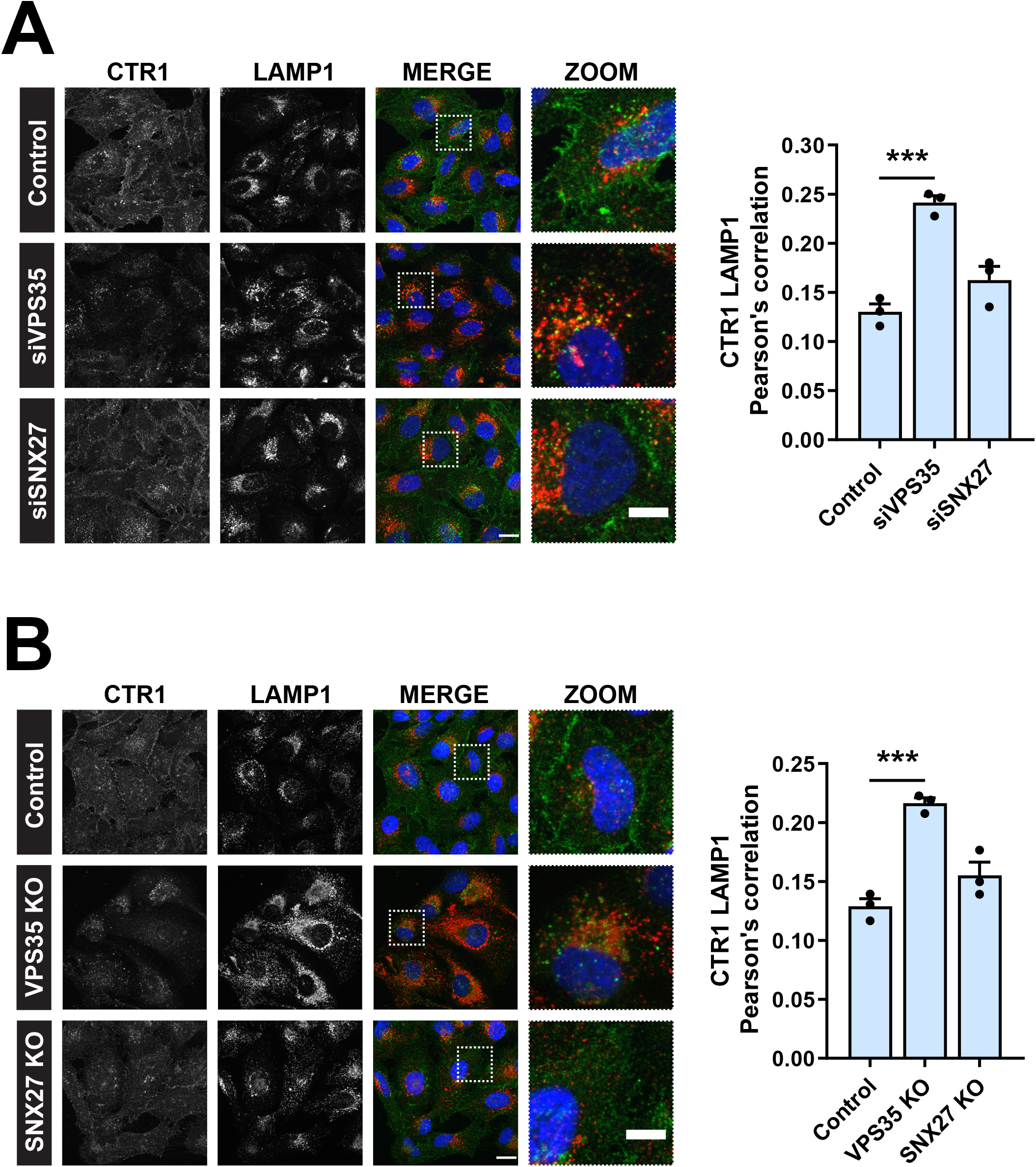
CTR1 is reduced from the cell surface and mis-sorted into lysosomes in the absence of VPS35, but not SNX27. (A) Antibody uptake with an antibody against endogenous CTR1. Immunofluorescence staining and colocalization analysis following a 2 hour incubation in CuCl_2_ (200 μM) at 37°C. Internalised CTR1 (green) and lysosomal marker LAMP1 (red) in control, VPS35− and SNX27-siRNA and (B) CRISPR-Cas9 knock out HeLa cells. The graphs represent means ± SEM; *n* = 3 independent experiments with > 100 cells per condition; one-way ANOVA followed by Dunnett’s multiple comparison. Scale bar, 20 μm and 10 μm in merge and zoom panels, respectively (A and B). \*\*\**P* < 0.001.

To ensure that the observed mis-sorting and reduction of CTR1 from the cell surface was specific to loss of retromer, we re-expressed VPS35 in VPS35 KO cells and examined the surface levels of CTR1 using flow cytometry. Upon re-expression of VPS35 the cell surface levels of CTR1 were restored to control levels (Figure 3A). This recovery of CTR1 levels at the cell surface following re-expression of VPS35 was also observed using surface biotinylation (Figure 3B). Together these data demonstrate that internalised CTR1 is retrieved from entering lysosomes and promoted for recycling back to the cell surface through a retromer-dependent pathway. Retromer is therefore required for controlling the cell surface residency of CTR1.

**Figure 3:**
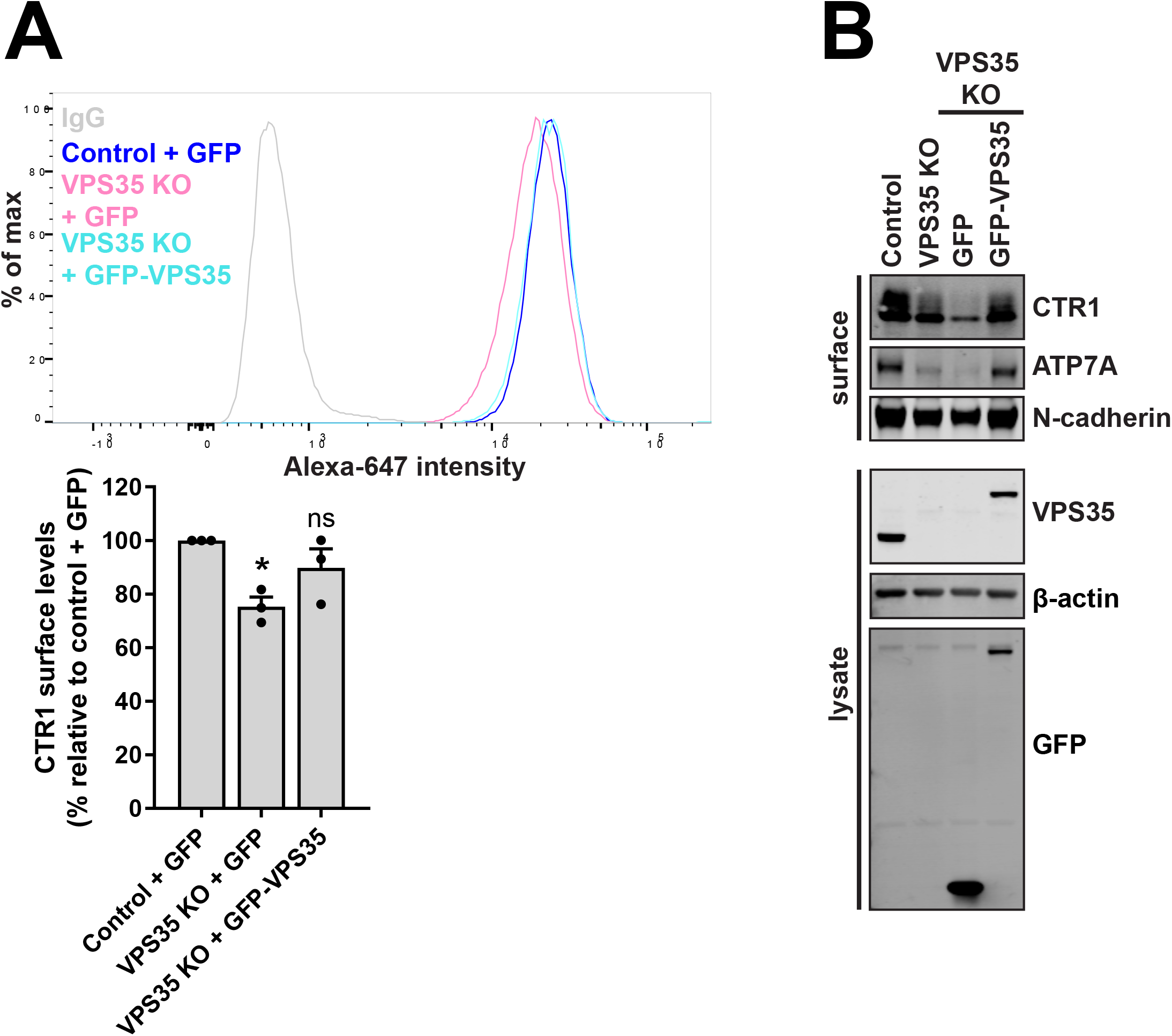
Re-expression of VPS35 rescues cell surface levels of CTR1. (A) Cell surface CTR1 levels were analysed by flow cytometry in control and VPS35 KO HeLa cells transduced with GFP control or GFP-VPS35 as indicated. Quantification represents the change in CTR1 cell surface levels relative to the control cells expressing GFP. The graphs represent means ± SEM; *n* = 3 independent experiments; one-way ANOVA followed by Dunnett’s multiple comparison. (B) Rescue of CTR1 cell surface levels assessed by surface biotinylation. The indicated HeLa cells were surface biotinylated and streptavidin agarose used to capture biotinylated membrane proteins. Surface abundance of the indicated proteins were examined by immunoblotting. \**P* < 0.05.

### Internalized CTR1 resides on retromer positive endosomes

Following a 30 minute exposure to extracellular copper, internalised CTR1 has been reported to localise to EEA1 and transferrin receptor positive early endosomes (Clifford et al., 2016). Therefore, we next sought to visualise whether following its copper-induced internalisation CTR1 traffics via a retromer-positive endosomal route. Cells were surface labelled on ice with an endogenous antibody to CTR1 and subsequently incubated at 37°C in excess copper-containing media (200 μM CuCl_2_) or standard growth media for 30 minutes prior to immunofluorescence processing. Cells were co-labelled with antibodies against endogenous CTR1 (red) and VPS35 (green). As anticipated, in cells maintained in standard low copper growth media CTR1 remained at the cell surface. Whereas, in cells where CTR1 had undergone copper-induced internalisation we found CTR1 extensively localised to VPS35-positive endosomes. Quantification using Pearson’s correlation determined a significant increase in the colocalization between CTR1 and VPS35 in cells exposed to copper (Figure 4A).

**Figure 4:**
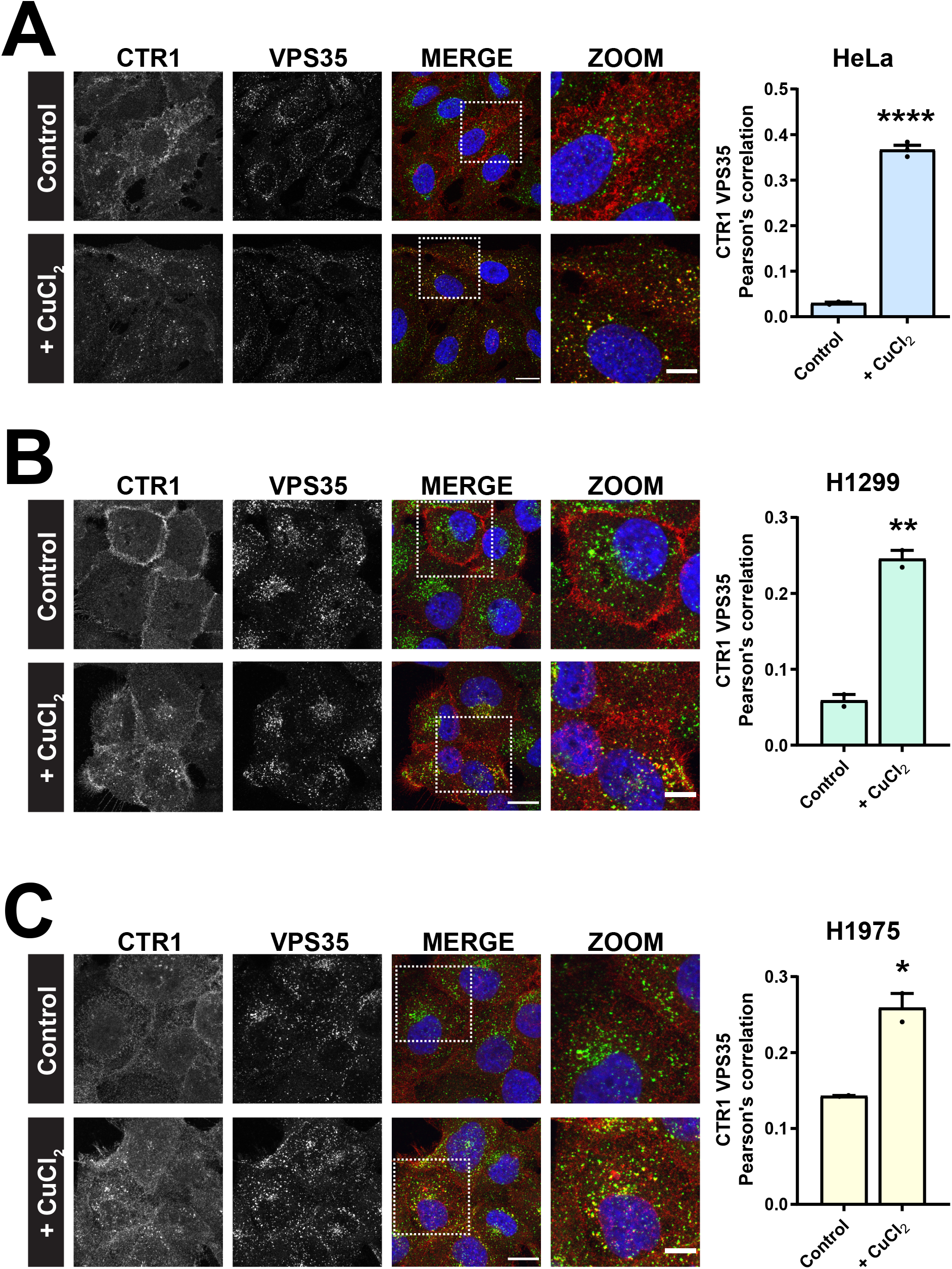
Following copper-induced internalization CTR1 colocalizes with retromer. (A - C) Antibody uptake experiments with an antibody against endogenous CTR1 in HeLa (A), H1299 (B), or H1975 cells (C). After 30 min incubation at 37°C in copper-containing media (200 μM CuCl_2_) internalised CTR1 (red) colocalized with endogenous VPS35 (green) on vesicular structures. In cells incubated in control media CTR1 remained extensively at the cell surface. Bars represent mean ± SEM from 3 independent experiments; Student’s *t* test (unpaired). Scale bar, 20 μm and 10 μm in merge and zoom panels, respectively (A, B and C). \**P* < 0.05; \*\**P* < 0.01; \*\*\*\**P* < 0.0001.

To determine whether this is a ubiquitous trafficking route for CTR1 we performed this CTR1 copper-induced internalisation assay in two different cell lines, H1299 and H1975 cells, both human non-small-cell lung cancer (NSCLC) cell lines. Notably, CTR1 was readily internalised in response to elevated copper in both these cell lines. As observed in HeLa cells (Figure 4A), in copper treated H1299 and H1975 cells internalised CTR1 localised to VPS35-positive endosomes and this correlated with a significant increase in the Pearson’s correlation between CTR1 and VPS35 (Figure 4 B and C). Together, these data provide further supporting evidence that in response to elevated extracellular copper internalised CTR1 is transported through retromer-positive endosomes.

### Retromer-dependent CTR1 trafficking is vital for copper homeostasis

Cellular adaptation to the ever-changing extracellular environment is fundamental to cell homeostasis and survival. Through regulating the constant flux of transmembrane proteins, the endosomal network is fundamental to the remodelling of the cell surface in response to extracellular cues (Curnock et al., 2019). To examine the importance of retromer-dependent CTR1 trafficking for copper homeostasis, we established a protocol to periodically incubate cells for 1 hour in high copper containing media (200 μM CuCl_2_) followed by a 1 hour incubation in normal growth media to effect a copper washout (Figure 5). This revealed a tight reciprocal coupling between CTR1 expression at the cell surface and the presence of elevated copper (Figure 5). Thus, upon addition of high copper CTR1 underwent endocytosis and underwent recycling back to the cell surface following copper washout. Entirely consistent with retromer regulating the recycling of internalised CTR1, the ability of cells to adapt to periodic exposure to elevated copper was completely lost upon retromer depletion (Figure 5).

**Figure 5:**
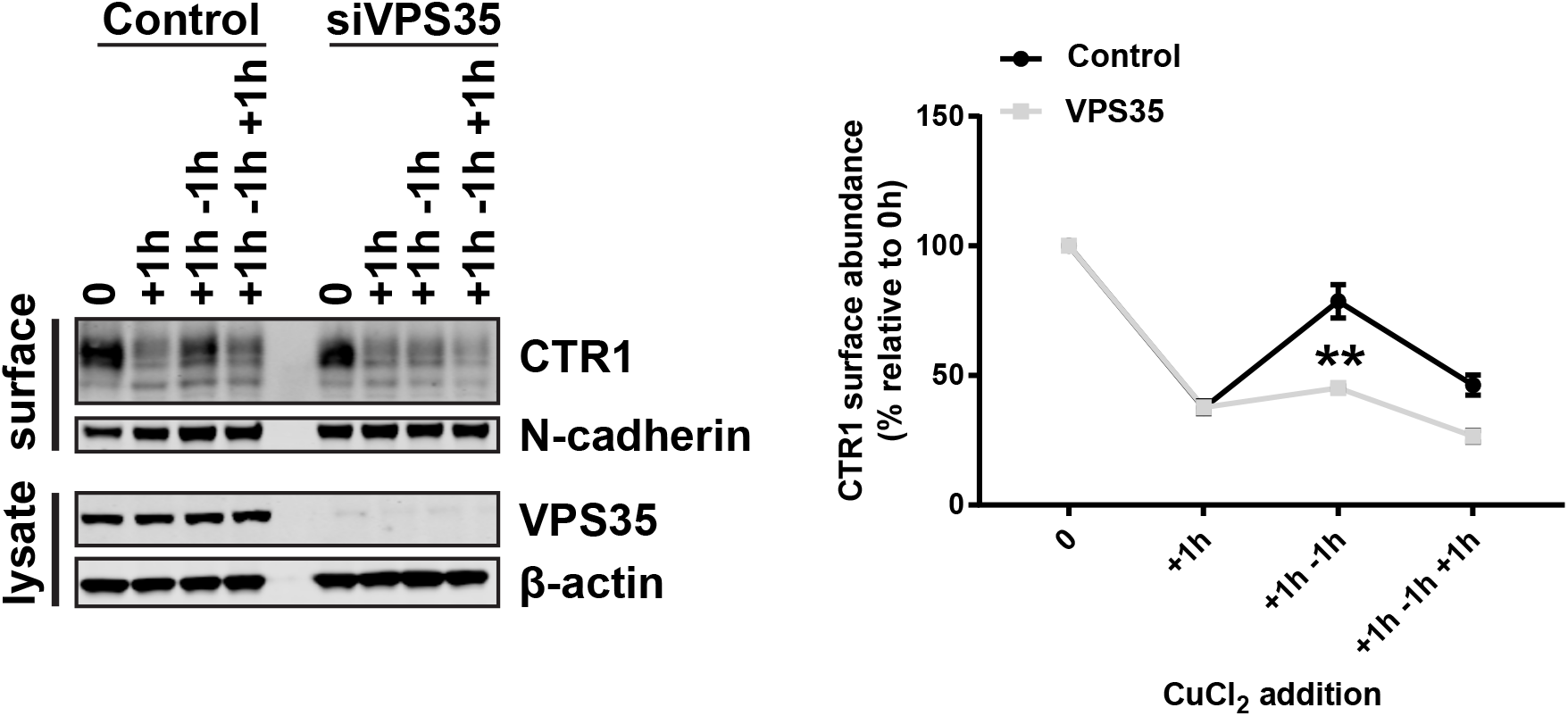
Retromer recycles CTR1 back to the cell surface in response to a reduction in extracellular copper concentration. CTR1 surface levels were analysed under copper-induced (200 μM CuCl_2_ for 1 hour) or copper washout (recovery in control media for 1 hour) conditions in control and VPS35-depleted cells. At the indicated timepoints cells were surface biotinylated and streptavidin agarose used to capture biotinylated membrane proteins. The surface abundance of CTR1 and the loading control N-cadherin were examined by immunoblotting. The graph represents the means ± SEM; *n* = 3 independent experiments; Student’s *t* test (unpaired). \*\**P* < 0.01.

To appreciate the role of retromer-dependent CTR1 trafficking in copper homeostasis we next performed MTT assays to assess the effect of increasing extracellular copper on metabolic activity as an indicator of cell viability. This colorimetric assay measures the ability of NAD(P)H-dependent oxidoreductases and dehydrogenases localized to cytoplasm and mitochondria to reduce MTT, a water-soluble tetrazolium dye, to a purple insoluble formazan (Berridge et al., 2005). Control and VPS35 KO HeLa cells were subjected to nine extracellular copper (CuCl_2_) concentrations from 3.125 μM to 800 μM for 48 or 72 hours and MTT reduction was measured by absorbance at 595 nm. In control cells after exposure to 400 μM copper for 48 hours a dramatic decline in metabolised MTT correlating with a decrease in cell viability was observed (Figure 6A). Strikingly, compared to control cells, VPS35 KO cells showed significantly reduced sensitivity to higher concentrations of extracellular copper. After 48 hours exposure to 400 μM copper, VPS35 KO cells still maintained the ability to metabolise MTT (Figure 6A). With prolonged exposure to copper for 72 hours, this difference in copper sensitivity between control and VPS35 KO cells became more pronounced (Figure 6B). After 72 hours, the toxic effects of the copper exposure were observed at lower concentrations in control cells however this was not as evident in VPS35 KO cells (Figure 6B). Importantly, we found that sensitivity to copper was restored upon re-expression of VPS35, thus demonstrating that the ability of the cell to sense and respond to toxic levels of extracellular copper relies on retromer function (Figure 6C).

**Figure 6:**
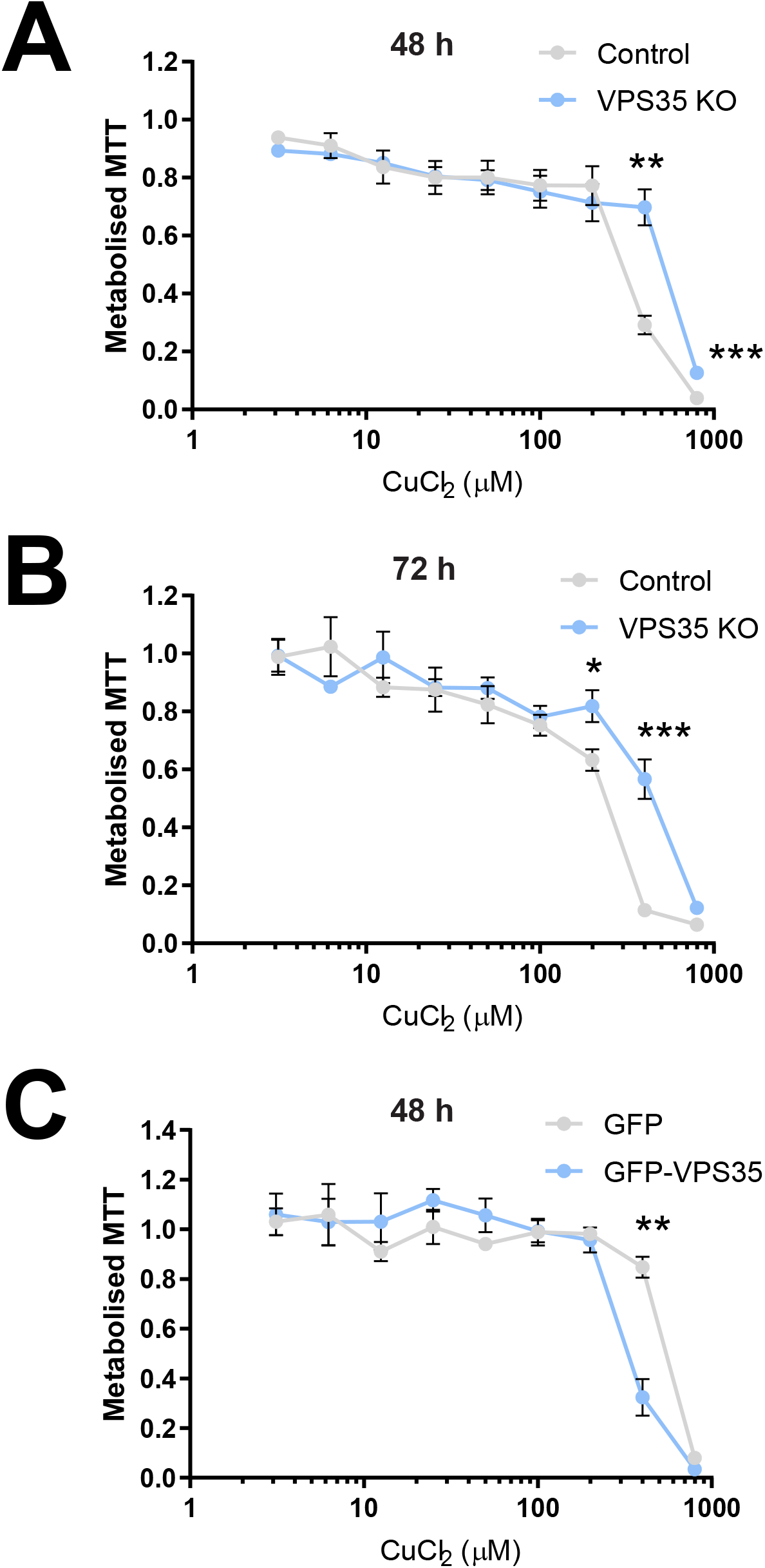
Retromer-dependent cargo recycling is necessary for copper homeostasis. (A - B) Control and VPS35 KO HeLa cells were incubated with nine concentrations of CuCl_2_ ranging from 3.12 to 800 μM for (A) 48 or (B) 72 hours. Each concentration was performed in quadruplicate. MTT reduction was measured by absorbance at 595 nm. Each datapoint represents the mean absorbance at 595 nm ± SEM, normalised to the baseline reading without addition of CuCl_2_; *n* = 4 independent experiments; Student’s *t* test (unpaired). (C) MTT assays were performed as described above subjecting VPS35 KO HeLa cells expressing GFP or GFP-VPS35 to increasing concentrations of CuCl_2_ for 48 hours. Each datapoint represents the mean absorbance at 595 nm ± SEM, normalised to the baseline reading without addition of CuCl_2_; *n* = 3 independent experiments; Student’s *t* test (unpaired). \**P* < 0.05; \*\**P* < 0.01; \*\*\**P* < 0.001.

### Retromer expression is required for maximal cisplatin toxicity

In addition to its role as the main transporter responsible for the cellular influx of copper, CTR1 has also been linked to the intracellular accumulation of platinum and therefore also platinum-containing cancer drugs (Howell et al., 2010). The efficiency of anti-tumour platinum-based drugs is directly related to the concentration of the drug that enters the cell. There is evidence that CTR1-dependent uptake controls the cytotoxicity of the widely used chemotherapeutic drugs cisplatin, carboplatin, and oxaliplatin (Holzer et al., 2006). Studies examining ovarian cancer cell lines and NSCLC patients samples identified that changes in CTR1 expression were linked to acquired cisplatin resistance and poor clinical outcome, whilst knockout of CTR1 in a murine model was shown to eliminate the responsiveness of tumours to cisplatin (Holzer et al., 2004; Jandial et al., 2009; Kim et al., 2014; Larson et al., 2009).

We have demonstrated that CTR1 surface abundance relies on retromer expression and knockout of VPS35 leads to a reduction in cellular sensitivity to extracellular copper. Therefore, given the evidence suggesting a dependency on CTR1 for cisplatin uptake and sensitivity, we next examined whether perturbation of retromer function affected cell viability in response to cisplatin exposure. We incubated control and VPS35 KO HeLa cells with nine increasing concentrations of cisplatin (from 0.39 μM to 100 μM). After 48 hours, a significant disparity in the sensitivity of control and VPS35 KO cells to cisplatin concentrations above 25 μM was observed (Figure 7A). These MTT assays revealed that in the absence of retromer, HeLa cells were partially desensitised to the toxic effects of cisplatin. To exclude that this effect was unique to HeLa cells, we also examined the human NSCLC cell line H1975. We reasoned that this was a suitable cell line to test this hypothesis as platinum-based chemotherapy drugs such as cisplatin and carboplatin form part of standard therapies commonly used to treat NSCLC (Cosaert and Quoix, 2002). Furthermore, our immunofluorescence analysis demonstrated that internalised CTR1 localised to VPS35-positive endosomes (Figure 4C) indicating that CTR1 undergoes an endosomal trafficking route in these cells.

**Figure 7:**
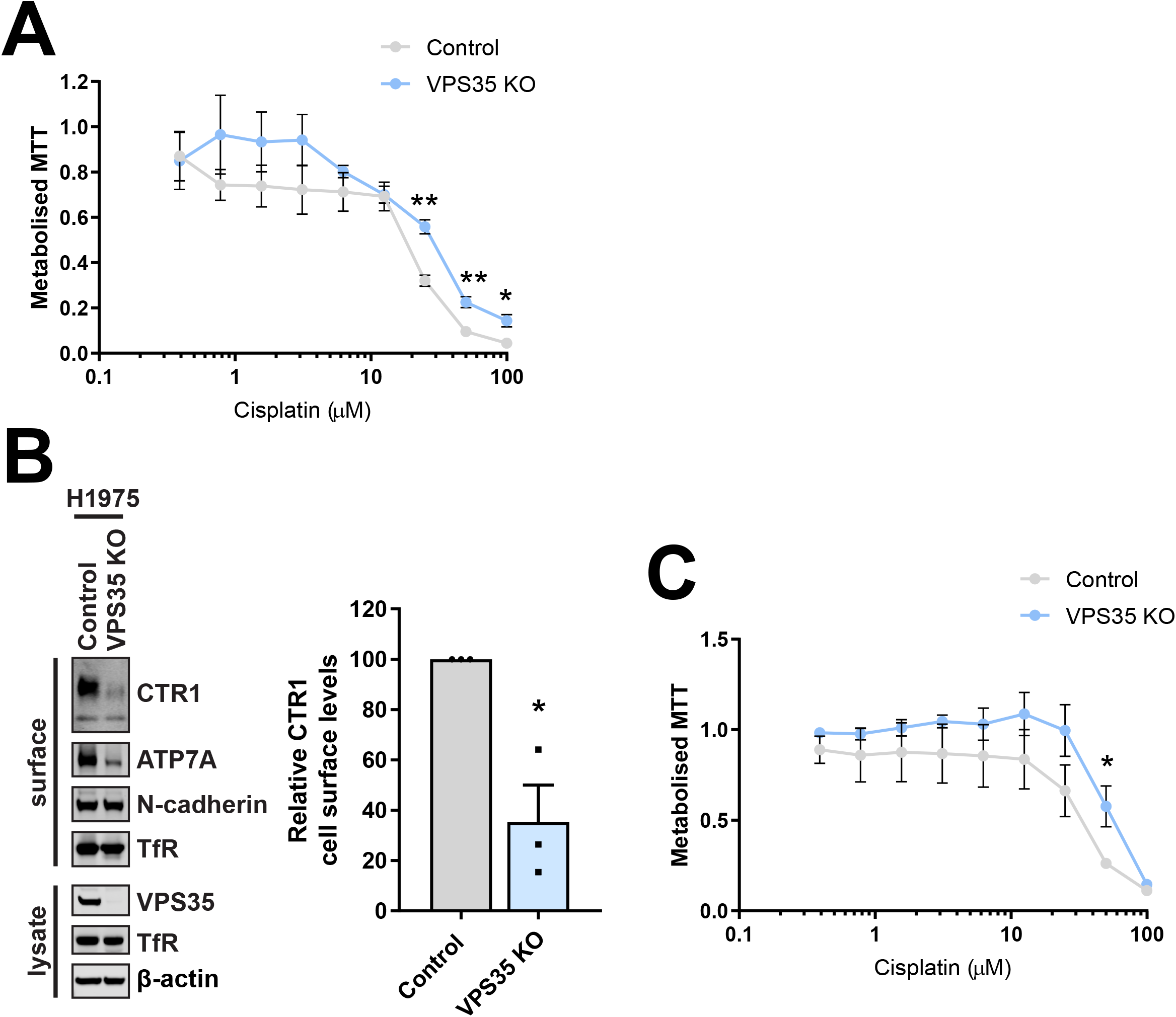
Retromer deficient cells show reduced sensitivity to the platinum-based drug, cisplatin. (A) Control and VPS35 KO HeLa cells were incubated with nine concentrations of cisplatin (from 0.39 μM to 100 μM) for 48 hours and MTT reduction was measured by absorbance at 595 nm. Each datapoint represents the mean absorbance at 595 nm ± SEM, normalised to the baseline reading without addition of cisplatin; *n* = 3 independent experiments; Student’s *t* test (unpaired). (B) CTR1 cell surface levels were assessed by surface biotinylation in VPS35 KO H1975 cells. Cells were surface biotinylated and streptavidin agarose used to capture biotinylated membrane proteins. Surface abundance of the indicated proteins were examined by immunoblotting. The quantification shows the mean ± SEM; *n* = 3 independent experiments; Student’s *t* test (unpaired). (C) Control and VPS35 KO H1975 cells were incubated with nine concentrations of cisplatin (from 0.39 μM to 100 μM) for 48 hours and MTT reduction was measure as described above; *n* = 4 independent experiments; Student’s *t* test (unpaired). \**P* < 0.05; \*\**P* < 0.01.

To first validate that retromer is required for CTR1 cell surface residency in this NSCLC cell line, surface biotinylation was performed following population based CRISPR-Cas9-mediated knockout of VPS35. Consistent with HeLa cells (Figure 1A and 3B), a significant reduction in the cell surface levels of CTR1 was observed in VPS35-depleted H1975 cells (Figure 7B). Notably, a loss of the copper transporting ATPase, ATP7A was also detected indicating that analogous to HeLa cells, in H1975 cells both the major cellular copper transporters responsible for maintaining copper homeostasis are affected by loss of retromer function. We next assessed using MTT assays whether perturbed retromer function effected sensitivity to cisplatin as observed in HeLa cells (Figure 5A). Following 48 hours in the presence of cisplatin, VPS35-depleted H1975 cells displayed a trend towards reduced sensitivity to cisplatin compared to the control counterparts (Figure 7C). Across different cell lines retromer is therefore required for cisplatin toxicity, implicating that disruption of retromer function may lead to a reduction in efficacy of platinum-based cancer drugs.

## DISCUSSION

Regulated copper homeostasis is fundamental in mammalian cells. Copper is an essential substrate for cell growth and differentiation, a cofactor of many enzymes and even recently identified as necessary for ULK1/2 signalling linking it to autophagy activity (Kim et al., 2008; Tsang et al., 2020). Therefore, it is unsurprising there are severe adverse effects associated with copper dyshomeostasis. Treatment for diseases resulting in acute cellular copper imbalances, such as Menkes disease are limited and prognosis is poor (de Bie et al., 2007). As the only known mammalian importer of copper, CTR1 could be a prospective pharmacological target. Therefore, gaining better insight into the membrane trafficking of CTR1 will enhance our mechanistic understanding of the regulation of cellular copper import and distribution, leading to a better appreciation for how this may be manipulated in disease.

The endosomal system is fundamental to organising the spatial distribution of integral membrane proteins such as metal transporters (Cullen and Steinberg, 2018). In response to elevated extracellular copper, ATP7A traffics from the TGN to the cell surface where it extrudes excess copper out of the cell (Petris et al., 1996; Polishchuk and Lutsenko, 2013). Maintaining the cell surface level of ATP7A, under basal conditions and following copper elevation, is reliant upon retromer and SNX27 orchestrating the endosomal retrieval and recycling of the internalised pump (Clairfeuille et al., 2016; Steinberg et al., 2013). This acute regulatory trafficking response requires a direct interaction between the PDZ binding motif present in the carboxy-terminal tail region of ATP7A and the PDZ-domain-containing SNX27 to mediate the endosomal delivery of ATP7A to the cell surface (Steinberg et al., 2013). In the present work we demonstrate that CTR1 is an additional retromer cargo and that retromer is required for the cell to adapt to changes in extracellular copper concentration. In response to elevated extracellular copper clathrin-dependent endocytosis of CTR1 is triggered, this serves to lower the CTR1 levels at the plasma membrane and restrict copper entry (Molloy and Kaplan, 2009). This cellular adaptation is vital to protect the cell from excessive copper accumulation. Here we demonstrate that retromer is required to facilitate the recycling of CTR1 back to the cell surface when extracellular copper levels are reduced (Figure 5). This acute recovery of CTR1 plasma membrane levels suggests that, in the presence of excess extracellular copper, the internal pool of CTR1 is not readily degraded and remains endosomally associated and accessible for retromer-mediated recycling (Figure 4A-C and Figure 8). We were unable to biochemically detect a direct interaction between CTR1 and retromer or retromer-associated adaptors (data not shown). We speculate therefore that interactions between CTR1 and retromer or retromer-associated endosomal recycling machinery are likely to be very transient and possibly even copper-dependent. It is interesting to postulate a system where copper modulates the conformation of CTR1 or even possibly key recycling machinery proteins to regulate their binding and dissociation, thereby coupling CTR1 recycling to extracellular copper levels.

**Figure 8:**
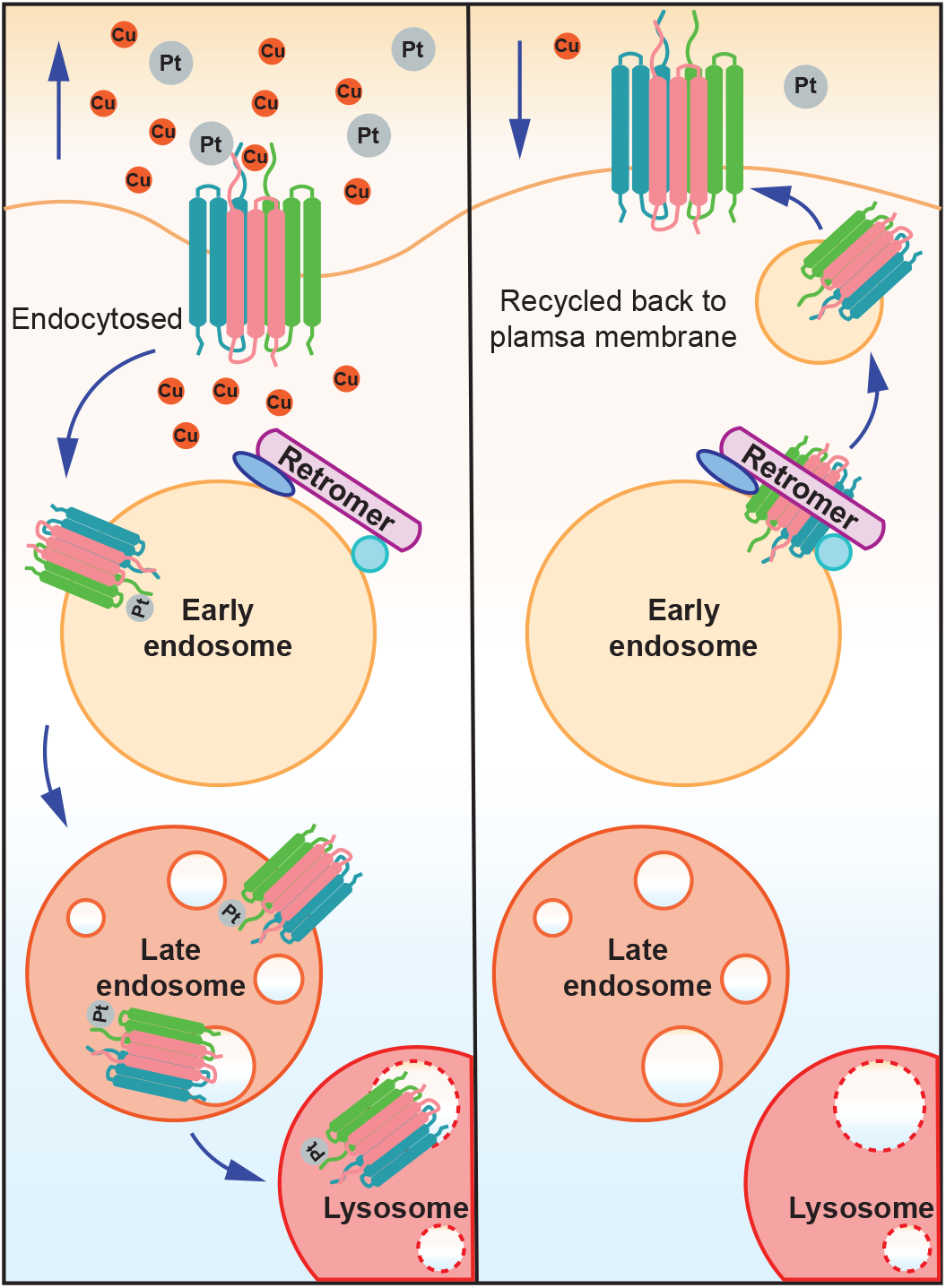
Model of retromer-dependent CTR1 recycling in response to changes in the concentration of extracellular copper or exposure to platinum-based drugs. In response to elevated copper or exposure to platinum-based drugs, such as cisplatin, CTR1 is endocytosed from the plasma membrane and sequestered on retromer-positive endosomes. In prolonged high copper or cisplatin conditions, CTR1 remains associated with the maturing endosome. Upon removal of extracellular copper or cisplatin, CTR1 is recycled back to the plasma membrane in a retromer-dependent manner.

Following the import of copper by CTR1, copper is transferred to the cytoplasmic copper chaperone ATOX1 (Kahra et al., 2016). CTR1 has a short 13-residue cytoplasmic C-terminal tail containing a highly conserved copper binding HCH motif that is necessary for relaying copper to the ATOX1 for delivery to copper-dependent enzymes (Kahra et al., 2016). NMR spectroscopy performed by Kahra et al., demonstrated that ATOX1 and the CTR1 tail peptide containing the HCH motif do not interact in the absence of copper, but copper exchange readily occurs from the copper loaded tail peptide to ATOX1. Copper transfer to high-affinity chaperones is therefore tightly controlled (Kahra et al., 2016).

For delivery of copper to the secretory pathway, the cytoplasmic copper chaperone, ATOX1, also interacts with the ATPases, ATP7A and ATP7B in a copper-dependent manner (Hamza et al., 1999; Larin et al., 1999; Pufahl et al., 1997). The copper-ATPases are known to form other copper-dependent protein interactions, such as with the antioxidant enzyme glutaredoxin 1 and the dynactin subunit p62 (Lim et al., 2006a; Lim et al., 2006b). Phosphorylation of the ATPases have also been identified to provide another level of regulation within the copper transport system with evidence of copper-stimulated phosphorylation and hyperphosphorylation (Vanderwerf et al., 2001; Voskoboinik et al., 2003). Understanding determinants that regulate copper-induced trafficking for CTR1 and the ATPases could provide fundamental information on how to modulate this pathway in disease.

It is established that highly conserved methionine residues within the import pore formed by the CTR1 trimer are essential for copper binding and uptake activity (Eisses and Kaplan, 2005; Puig et al., 2002; Ren et al., 2019). Interestingly, the pore diameter does not permit permeation of larger cisplatin molecules (Ren et al., 2019). Cisplatin has been proposed to bind to an extracellular proportion of CTR1 and distinct from copper, enter the cell via endocytosis (Larson et al., 2010). Despite their differences in entry mechanism, the density of CTR1 at the cell surface is pertinent to the amount of both copper and cisplatin capable of entering the cell.

Indeed, consistent with a reduction in CTR1 at the cell surface correlating with lower copper and cisplatin uptake, retromer deficient cells were significantly less sensitive to copper loading and cisplatin exposure (Figure 6A-C and Figure 7A and C). These findings suggest a link between retromer-dependent CTR1 recycling and sensitivity to these exogenous cellular stressors. Cisplatin resistance has been linked to CTR1 degradation and knockout in cells and *in vivo* (Holzer et al., 2006; Larson et al., 2009). Furthermore, resistance to platinum-based cancer drugs is a significant obstacle in the effectiveness of therapy. Low expression of CTR1 has been associated with poor clinical outcome in NSCLC patients who had received platinum-based chemotherapy (Kim et al., 2014). Interestingly, the copper transporting ATPases, ATP7A and ATP7B, have been suggested to chelate and/or promote the cellular efflux of platinum-based drugs (Dolgova et al., 2009; Gupta and Lutsenko, 2009; Katano et al., 2004; Polishchuk and Polishchuk, 2016; Safaei and Howell, 2005). Studies have revealed a correlation between elevated ATP7B expression and reduced effectiveness of cisplatin chemotherapy in cancer patients (Miyashita et al., 2003; Nakayama et al., 2004). Consequently, targeted downregulation of ATP7B expression and trafficking is currently being explored as a potential strategy to increase the efficacy of cisplatin therapy (Mangala et al., 2009; Mariniello et al., 2020).

Overall, through a variety of approaches we have shown that retromer is required for the cell surface localisation and copper-dependent recycling of CTR1 (Figure 1A, C and Figure 3A, B and Figure 7B). We propose a working model where upon copper- or cisplatin-dependent internalisation CTR1 enters retromer positive endosomes. Here CTR1 can undergo one of two fates: in sustained elevated copper CTR1 remains associated with the maturing endosome and is degraded within the lysosome (Guo et al., 2004; Liu et al., 2007; Petris et al., 2003) or, as we demonstrate in the present work, following a reduction in extracellular copper levels CTR1 becomes available for retromer-dependent recycling back to the cell surface (Figure 8). Exploring this model further will reveal greater insight into the regulation and trafficking of CTR1 and may suggest approaches to manipulate cellular copper import and distribution in disease and modulating cellular sensitivity and toxicity to platinum-based treatments.

## MATERIALS AND METHODS

### Antibodies

Antibodies used in the study were: (WB: western blot, IF: immunofluorescence, FC: Flow cytometry): Mouse monoclonal antibodies raised against SNX27 antibody (clone 1C6, Abcam, Ab77799, WB), GFP (clones 7.1/13.1, Roche, 11814460001, WB), LAMP1 (clone H4A3, Developmental Studies Hybridoma Bank, IF), N-Cadherin (clone 32, BD Biosciences, 610920, WB), ATP7A (clone D9, Santa Cruz sc-376467, WB), Transferrin receptor (clone H68.4, Invitrogen 13-6890, WB), β-actin (Sigma-Aldrich, A1978, WB). Rabbit monoclonal antibodies raised against VPS35 (Abcam, ab157220, WB), SLC31A1/CTR1 (Abcam, ab129067 WB/IF/FC). Rabbit polyclonal antibodies raised against VPS35 (Abcam 97545, IF), Isotype Ctrl (clone Poly29108, Biolegend 910801, FC). For Odyssey detection of western blots, the following secondary antibodies were used; goat anti-mouse 680 (Invitrogen), goat anti-rabbit 800 (Invitrogen).

### Cell culture conditions

All cell lines were maintained at 37°C with 5% CO_2_ atmosphere. HeLa cells were cultured in Dulbecco’s Modified Eagle Medium containing 4.5 g/L glucose (DMEM) (D5796; Sigma-Aldrich), supplemented with 10% (v/v) fetal bovine serum (FBS). H1975 cells were maintained in RPMI 1640 medium (R0883; Sigma-Aldrich), supplemented with 10% (v/v) FBS and 2mM L-glutamine. For copper supplement experiments cells were incubated for the indicated time at 37°C in medium containing CuCl_2_ (Sigma-Aldrich).

### Transfections

DNA was transiently transfected into cells using FuGENE 6 transfection reagent (Promega), according to the manufacturer’s instructions. The gRNAs for CRISPR genome editing were cloned into the CRISPR-Cas9 plasmid px330. The gRNAs used in this study were VPS35, 5′-GTGGTGTGCAACATCCCTTG-3′ and SNX27, 5’-GGCTACGGCTTCAACGTGCG-3’. CRISPR-Cas9 plasmids were cotransfected with a puromycin resistance–expressing plasmid, and cells were subjected to puromycin selection 24 h later.

For siRNA-based knockdown, cells were reverse-transfected using DharmaFECT 1 (GE Healthcare) and then transfected again 24 h later according to the manufacturer’s instructions. 48 h after the second transfection, cells were lysed or fixed and processed for immunofluorescence. For VPS35 suppression, a combination of oligonucleotides 3 (sequence 5′-GUUGUUAUGUGCUUAGUA-3′) and 4 (sequence 5′-AAAUACCACUUGACACUUA-3′) of the Dharmacon ON-TARGETplus VPS35 siRNA SMARTpool was used. For SNX27 suppression, Dharmacon ON-TARGETplus SNX27 siRNA SMARTpool (cat. #L-017346-01) was used. The non-target control siRNA used was Dharmacon ON-TARGETplus non-targeting siRNA no. 2 (cat. #D-001810-02).

### Quantitative Western blot analysis

For Western blotting, cells were lysed in PBS containing 2% Triton X-100 and Pierce protease and phosphatase inhibitor tablets (Thermo Fisher Scientific). The protein concentration was determined with a BCA assay kit (Thermo Fisher Scientific), and equal amounts were resolved on NuPAGE 4–12% precast gels (Invitrogen). Proteins were transferred from gels to PVDF membrane (Immobilon-FL, pore size 0.45 μm, Millipore, catalogue number IPFL00010) for immunoblotting. Transfer was performed in transfer buffer (25 mM Tris, 192 mM glycine, 10% methanol) at 100 V for 70 minutes. Membranes were blocked in 5% milk in PBS-T (PBS containing 0.1% Tween 20) prior to incubation with the appropriate primary antibodies and fluorescently labelled secondary antibodies. Detection and quantification were carried out using an Odyssey infrared scanning system (LI-COR Biosciences).

### Cell surface biotinylation

Cells were surface biotinylated 72 h post siRNA transfection or after the indicated treatment with membrane-impermeable biotin (Thermo) at 4 °C to prevent endocytosis. Post-biotinylation, cells were lysed in PBS containing 2% Triton X-100 and Pierce protease and phosphatase inhibitor EDTA-free tablets, pH 7.5 and lysates were cleared by centrifugation. Equal amounts of protein from the control, indicated knockdown, knockout or treatment lysate were then added to streptavidin Sepharose to capture biotinylated proteins. Streptavidin beads and lysates were incubated for 30 mins at 4 °C before washing in PBS containing 1.2 M NaCl and 1% Triton X-100. Proteins were eluted in 2× NuPAGE LDS Sample Buffer (Life Technologies) by boiling at 95 °C for 10 min, then separated by SDS– PAGE and subjected to quantitative Western blot analysis (as described above).

### Immunofluorescence

Cells were washed three times in PBS, fixed in 4% paraformaldehyde (PFA) and then washed in PBS and permeabilized with 0.1% Triton X-100 or saponin. Fixed cells were then blocked in PBS containing 1% BSA and incubated with primary antibodies and appropriate secondary antibodies (Alexa Fluor; Thermo Fisher Scientific) in PBS containing 0.1% BSA. PBS-washed coverslips were mounted onto glass slides with Mowiol (Sigma-Aldrich).

### Antibody uptake assays

For the CTR1 trafficking assays, cells were seeded onto sterile coverslips. The following day, cells were surface labelled by incubation with an antibody against CTR1 for 30 minutes on ice. Cells were then incubated at 37°C in excess copper-containing medium (200 μM CuCl_2_) or standard growth medium for either 30 minutes or 2 hours as indicated, at which point, cells were fixed in 4% PFA and processed for immunofluorescence staining as described above.

### Flow cytometry

Cells were washed three times in PBS (without Ca or Mg), and then incubated for 20 minutes with Accutase to detach cells. 200,000 resuspended cells were transferred to 5 ml round-bottom polystyrene tubes (BD Biosciences). Cells were reconstituted in PBS (without Ca or Mg), containing 1% BSA and incubated with rabbit monoclonal CTR1 antibody (ab129067) or an isotype control antibody (Biolegend, clone Poly29108) for 1 hour on ice. Cells were washed once in ice-cold PBS (without Ca or Mg) and incubated for 30 min on ice with Alexa Fluor 647 conjugated donkey anti-rabbit secondary antibody (Invitrogen, A31573) in PBS (without Ca or Mg), containing 5% BSA. The cells were washed again and resuspended in PBS (without Ca or Mg) containing 1% BSA and propidium iodide. Fluorescent signals were measured using a FACS CantoII-F60 machine (BD Biosciences, Oxford, UK). Data were analyzed using Flowjo 7.2.5 software (Flowjo, Ashland, OR).

### MTT assay

Cells were seeded into 96-well plates at a density of 8 × 10^3^ cells/well in DMEM without phenol-red supplemented with 10% (v/v) FBS. The following day, cells were treated with serial dilutions of copper (CuCl_2_ 3.125 μm to 800 μm) or cisplatin (0.39 μM to 100 μM). Each condition and an untreated control were performed in quadruplicate. After incubation at 37°C with 5% CO2 atmosphere for either 48 or 72 hours as indicated, 20 μl 5 mg/ml MTT (3-(4,5-Dimethylthiazol-2-yl)-2,5-Diphenyltetrazolium Bromide; Life Technologies M6496) was added to each well, and the plates were incubated for an additional 3.5 hours at 37°C. MTT was removed and 150 μl DMSO was added to each well and mixed thoroughly by pipetting. MTT absorbance was read at 595 nm using a microplate reader. The average of the quadruplicate reading for each condition was expressed as percentage of the untreated control.

### Image acquisition and analysis

Microscopy images were collected with a confocal laser-scanning microscope (SP5 AOBS; Leica Microsystems) attached to an inverted epifluorescence microscope (DMI6000; Thermo Fisher Scientific). A 63× 1.4 NA oil immersion objective (Plan Apochromat BL; Leica Biosystems) and the standard SP5 system acquisition software and detector were used. Images were captured using Application Suite AF software (version 2.7.3.9723; Leica Microsystems) and then analyzed with Volocity 6.3 software (PerkinElmer). For colocalization studies, Pearson’s correlation (measuring the correlation in the variation between two channels) was measured using Volocity 6.3 software (PerkinElmer).

### Statistical Analysis

All statistical analyses were performed using Prism 7 (GraphPad Software). For comparing two groups, statistical analysis was performed using student’s *t* test (unpaired). For multiple comparisons, one-way ANOVA followed by Dunnett’s multiple comparison test were used. Data distribution was assumed to be normal, but this was not formally tested. All quantified Western blot and confocal colocalization data are the mean of at least three independent experiments. Graphs represent means and SEM.

## Author contributions

RC conceived the study, performed all of the experiments, wrote the draft manuscript and made the figures. RC and PJC edited and approved the final manuscript.

